# Identification and isolation of migratory subpopulations of cancer cells using microfluidics with fluid walls

**DOI:** 10.64898/2026.11.11.765957

**Authors:** Joseph A. E. Morgan, Daniele T. Cotton, Srinivasa R. Rao, Peter R. Cook, James R. Edwards, Alfonso A. Castrejón-Pita, Claire M. Edwards, Edmond J. Walsh

**Author notes:** Corresponding authors: Claire M. Edwards, Edmond J. Walsh **Email:**. Contributed equally: Joseph A. E. Morgan, Daniele T. Cotton. **Author contributions:** J.A.E.M, D.T.C, S.R.R, C.M.E and E.J.W designed research, J.A.E.M and D.T.C performed research, all authors analyzed data, wrote and reviewed the paper.

## Abstract

Microfluidic assays have great potential for investigating cancer cell migration, a key process underlying metastasis which remains a major clinical challenge. However, adoption by biologists is limited due to complex fabrication and operation methods combined with difficulty in retrieving cells of interest. We jet-print dumbbell-shaped microfluidic environments with fluid walls in standard Petri dishes in seconds using standard growth media and immiscible oil. Prostate cancer cells and chemoattractant (bovine serum) are introduced into each end using local transient differences in Laplace pressure to enable spatial confinement. Automated live-cell imaging monitors chemotaxis towards serum using standard microscopes, as the confining fluid walls are optically transparent. Distinct subpopulations of migratory cells exhibit different morphologies and motilities, assessed using a custom machine-learning segmentation model. Then, we leverage the reconfigurability of fluid walls to isolate and retrieve rapidly-migrating single cells. This demonstrates that fluid-walled microfluidics is a versatile and accessible approach for investigating cancer cell migration at single-cell resolution, offering new opportunities to examine cellular heterogeneity in metastasis.

## Introduction

Cell migration is a critical step during metastasis. For example, in prostate cancer, cells shed from the primary tumour can invade surrounding tissue, enter blood vessels, and disseminate throughout the body (**Figure 1A**) [1]. A frequently cited study reports that 90% of men who die from prostate cancer also exhibit bone metastases [2]. Consequently, many *in vitro* migration assays have been developed, e.g., involving Transwells, Dunn chambers, wound-healing, and complex matrices [3–7]. However, existing methods have their respective limitations: for example, Transwell assays provide limited information on cell morphology during migration, and wound-healing (scratch) assays are generally unsuitable for studying chemotaxis [8].

**Figure 1.**
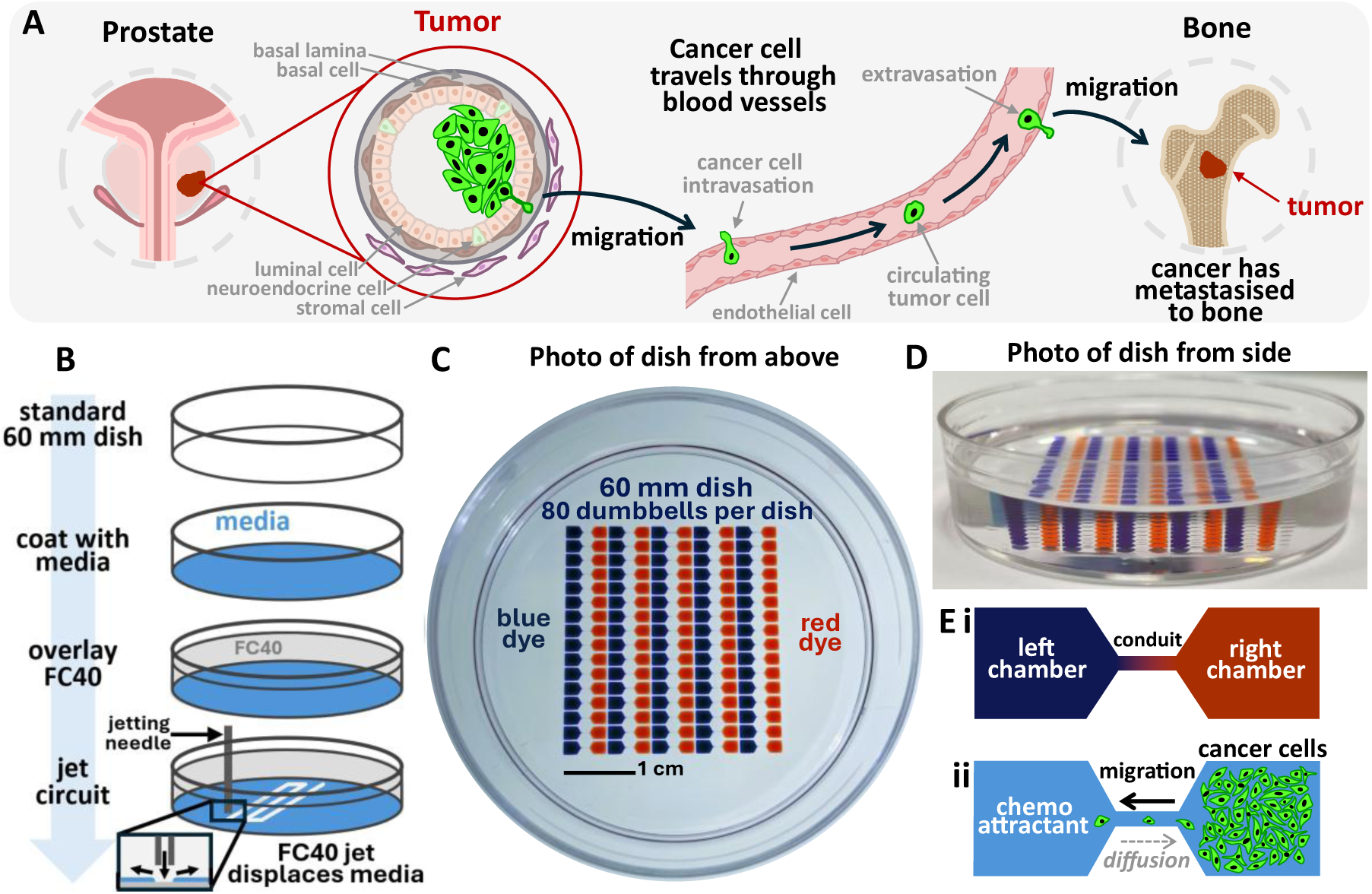
A microfluidic assay with fluid walls for cell migration. **(A)** Primary prostate tumors can metastasise to the bone via cell migration and intra-/extravasation through the bloodstream. **(B)** To study this process, we engineered fluid-walled microfluidic dumbbells. A thin film of culture medium covered a standard 60 mm dish, FC40 fluorocarbon was overlaid, and FC40 jetted through the medium to create fluid walls. **(C)** 80 dumbbells were fabricated on a single 60 mm dish. Image shows red and blue dyes in opposing chambers (top-down image acquired with an iPhone camera). **(D)** Side-on photo of dish. Dumbbells are pinned to the polystyrene surface under FC40. **(E)** Chambers can be addressed independently. For example, **(i)** dyes to aid circuit visualisation, or **(ii)** seeding cells in one chamber and introducing a chemo-attractant in the other which diffuses through the conduit, promoting migration from the right chamber to the left.

Microfluidic approaches have emerged as promising alternatives, especially for high-throughput assays, providing greater flexibility in assay design and enabling real-time monitoring of directed migration through microchannels [9], [10]. Such platforms allow precise control over cellular microenvironments and have been used to identify distinct migratory cancer cell subpopulations alongside non-migratory [11], [12], [13]. In some cases, microfluidic systems have facilitated the isolation and retrieval of migratory cells, enabling downstream comparisons that have demonstrated increased metastatic potential *in vivo* compared to non-migratory cells [13]. However, such approaches are typically limited to low-resolution classification (for example, migratory or non-migratory) as cells are pooled during retrieval, restricting single cell analysis to interrogate heterogeneity within migratory subpopulations. Furthermore, microfluidic devices are typically sealed within polydimethylsiloxane (PDMS), limiting access to cells within enclosed microchannels. Open-microfluidic systems partially address these limitations by improving accessibility [14], [15] but often rely on complex fabrication techniques that may present a barrier to adoption by biologists [16].

Microfluidics with fluid walls offers an effective solution where geometries are defined by immiscible fluids that form liquid interfaces rather than solid walls [17]. Spatially distinct culture systems can be formed using standard cell-culture media and confined by the immiscible and bio-inert fluorocarbon oil, FC40, with geometry maintained by interfacial forces. Such culture systems can be made in seconds on standard tissue-cultureware and, as confining walls are liquid and transparent, they are easily pierced with pipets so that all regions are fully accessible. Moreover, fluid walls can be reconfigured during assays to allow isolation and retrieval of cells of interest [18]. The advantages of using fluid-walled microfluidics in cell migration studies have been demonstrated in assays of macrophage and bacterial chemotaxis [19], [20].

We present a microfluidic assay with fluid walls to study the migration of PC3 prostate cancer cells, derived from a vertebral adenocarcinoma metastasis [21]. PC3 cells possess high metastatic potential in *in vivo* models and have been used previously in various migration assays [22], [23], [24]. We jet-print arrays of fluid walls shaped like dumbbells, each comprising two chambers connected by a narrow fluid conduit. Modulating local volumes directly alters height, but not footprint, and hence local pressures. In our system, geometries are sufficiently small that interfacial forces outweigh gravitational ones (Bond number < 1), and behaviour is governed by gradients in Laplace pressure; adding or withdrawing volume enables control over flow direction [25], [26]. We exploit pressure gradients to confine cells and chemo-attractant (foetal bovine serum, FBS [27]) to respective chambers [25]. Over 3 days, cells migrate through the conduit towards FBS. We then identify sub-populations of highly-migratory cells, which likely possess higher metastatic potential compared to non-migrating ones [12], [13]. We then reconfigure the fluid walls – a process unique to this method, to isolate the most migratory cell(s) from the majority and retrieve them.

The accessibility of machine-learning tools for cell segmentation is advancing rapidly [28]. We trained a custom model using Cellpose-SAM [29], a framework that combines the strong generalization of Cellpose with the Segment Anything Model [30] to automate tracking of individual cells and elucidate cell characteristics. This uncovered striking morphological heterogeneity at a single-cell level in migrating sub-populations. In summary, we demonstrate the versatility of microfluidics with fluid walls for studying single-cell migration and subsequently apply this platform to identify and characterize heterogeneity in sub-populations of migratory cancer cells.

## Results

### Jet-printing microfluidic dumbbells with fluid walls

Microfluidic dumbbells were fabricated on standard 60 mm polystyrene culture dishes (Corning, Corning, NY, USA) using a modified fluid printer (isoCell, iotaSciences Ltd, Oxford, UK). First, the dish was coated with a thin layer of medium (1 mL, withdrawn manually to leave a thin film) and overlaid with 3 mL of the immiscible fluorocarbon, FC40. A submerged microjet of FC40 was then used to displace the underlying medium, leaving behind stable FC40 fluid walls held in place by interfacial forces (**Figure 1B**). This process is referred to as jet-printing [31]. FC40 is bio-inert, gas permeable, and has refractive index close to water, making it well-suited to live-cell culture and imaging [32]. The jet path was controlled via the fluid printer, enabling rapid generation of any two-dimensional (2D) pattern. Using this approach, we produced an array of 80 dumbbells (5 columns, 16 rows) in ⁓ 30 s (**Figure 1C, D**). Each dumbbell consisted of two chambers (1.8 mm x 1.8 mm) connected by a narrow conduit (1 mm length, 0.2 mm width). Since dumbbells are confined by a fluid-fluid interface, all regions are fully accessible. After lowering a dispensing needle (connected to a syringe pump, controlled via the fluid printer) through the interface, media can be infused/withdrawn locally into/from each chamber independently (**Figure 1Ei**, see red and blue dye). This approach can be used to seed cells in one chamber and deliver chemo-attractant to the other (**Figure 1Eii**). Dumbbells were fabricated using medium + 0.5% FBS, which we found to be the minimum serum concentration required to ensure stable fluid walls (supplementary **Figure S1**).

### Controlling flow in microfluidics with fluid walls

Unlike conventional microfluidic systems bounded by solid walls, fluid walls – while tightly pinned to the underlying substrate – are free to increase or decrease in height when media is added or withdrawn. Consequently, regions within a dumbbell may have different radii of curvature depending on local volumes, and hence different pressure rises across the fluid interface – Laplace pressures – which are given by:

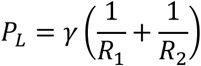

In the case of a small drop deposited on a surface, *PL* = 2*γ*⁄*R*, with *R* the radius of curvature of the drop and *γ* the interfacial tension (for DMEM + 10% FBS and FC40, *γ* = 22.8 mN/m [33]). By approximating chamber 3D geometry as that of a spherical cap, this can be expressed in terms of chamber heights ℎ and footprint radius *a*:

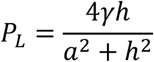

and height estimated from the volume *V* in the chamber [33]:

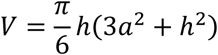

Consider a dumbbell circuit at equilibrium where both chambers have the same footprint geometry and equal volume with contact angles, *θ* less than 90⁰ (**Figure 2**). Adding volume to the left chamber will increase chamber height (and reduce the radius of curvature) causing an increase in Laplace pressure. Consequently, this induces flow from the left chamber to the right until pressures in each chamber equilibrate. These principles underpin our ability to exploit circuit dynamics to achieve spatial control within the circuit and create functional migration assays. For further detail regarding circuit pressure dynamics and 3D geometry, see supplementary **Figure S2**.

**Figure 2.**
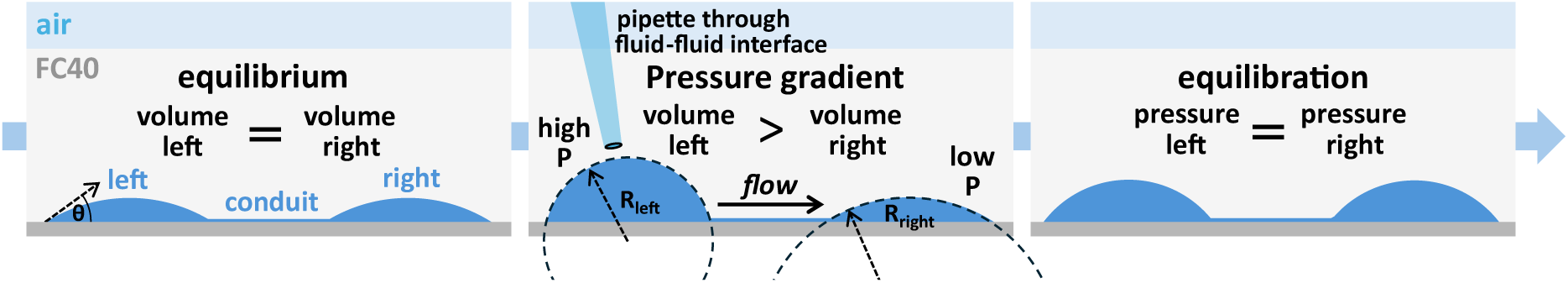
Controlling flow in microfluidics with fluid walls. Two chambers with the same footprint geometry and volume connected by a fluid conduit are at equilibrium. Adding volume to the left chamber causes a local increase in Laplace pressure. This pressure imbalance drives flow from the left to the right chamber until pressures equilibrate.

### Enabling spatial confinement of cells and chemo-attractant

Following jet printing, displaced medium accumulated within chambers such that each chamber contained **⁓** 150 nL (estimated by comparing dish weight before and after adding the thin film of medium and calculating μL/mm^2^ of medium displaced by the FC40 fluid walls). To standardize chamber volumes and remove residual serum, dumbbell chambers were washed with 600 nL phosphate-buffered saline (PBS) and 800 nL withdrawn. During washing, the estimated peak contact angle was **⁓** 33°. Withdrawing more than the chamber volume ensured near-complete removal of medium, while also extracting a small volume of FC40 overlay, thereby minimizing any variability in the volume of media in chambers. The efficacy of the PBS wash in removing residual serum is shown in supplementary **Figure S3,** where fluorescent bovine serum albumin, BSA, is used as a surrogate for serum. Following the wash, dumbbells were coated with 100 nL FNC coating mix (Athena Enzyme Systems, Baltimore, MD, USA) to promote cell attachment. Next 400 nL serum-free medium was added to the left chamber to generate a pressure gradient driving flow toward the right chamber. A smaller volume of cell suspension (100 nL, **⁓** 500 PC3 cells, transduced to express enhanced green fluorescent protein, eGFP) was then seeded into the right-hand chamber. The pressure difference between the two chambers ensured flow was from left to right, and this confined deposited cells to their chamber (**Figure 3A).** Individual cells are clearly visible in this chamber. Cells adhered within 3 h. As flow between chambers persisted well beyond this, cells remained in the right-hand chamber (see supplementary **Figure S4** for flow estimations and supplementary **Figure S5** for optimisation of cell seeding number).

**Figure 3.**
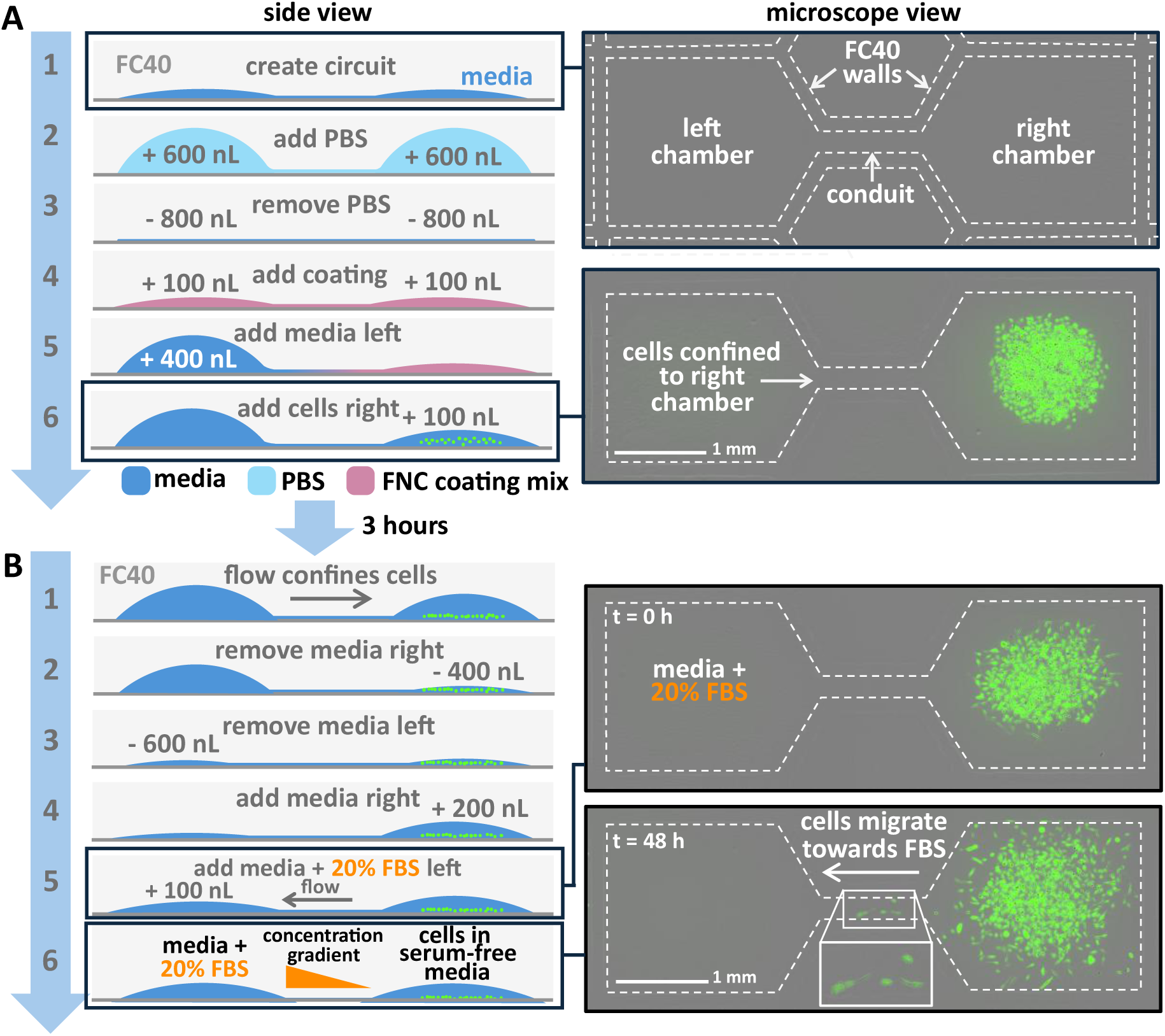
Workflow for introducing cells and chemo-attractant into dumbbells. Low-power fluorescence microscope views of the circuit at the stages indicated are shown on the right. **(A)** (1) Dumbbells were fabricated by jetting FC40 through a thin film of culture media to create fluid walls (image on right), (2,3) washed with PBS, and (4) coated with FNC coating mix. To confine cells, (5) 400 nL of medium was added to the left chamber to generate a gradient in Laplace pressure, driving flow from left to right. (6) PC3-eGFP cells were now added to the right-hand chamber (fluorescence image on right). **(B)** (1) Differences in Laplace pressure ensure flow is to the right (sustained for >3 hours), so cells are confined to the right. (2) After 3 h, medium is removed on the right, and (4) medium added to drive flow from right to left and hence initially confine chemo-attractant. (5) Chemo-attractant (media + 20% FBS) was introduced into the left. Differences in chamber volume were used to control flow and confine serum to the left chamber. Flow persisted for ⁓ 2 hours, after which (6) diffused through the conduit, establishing a stable diffusion gradient after ⁓ 3.5 hours, promoting cell migration to the left. Images on the right show cells migrating towards 20% FBS.

Once cells had adhered, the same principles were applied to introduce and confine a chemo-attractant into the left-hand chamber (**Figure 3B**). Here, 200 nL serum-free medium was added to the right-hand chamber containing cells, and 100 nL medium + 20% FBS was added to the left, thus generating flow from right to left (ensuring chemo-attractant was confined to the left-hand chamber). After ⁓ 2 hours (approximate equilibration time), serum then diffused into/through the conduit, estimated to establish a stable diffusion gradient after an additional ⁓ 3.5 hours (which was estimated to remain stable for the duration of the assay) and some cells began to migrate along the concentration gradient from right to left. See supplementary **Figure S6** for estimation of diffusion rates, and **Figure S7** for effect of serum concentration and chamber volume on migration.

### Cell migration through the conduit

Following addition of chemo-attractant, dumbbells were placed in a standard incubator (5% CO_2_, 37°C) and imaged hourly thereafter using an Incucyte live-cell imaging system (phase-contrast and fluorescence). The excellent optical properties of fluid walls enabled clear visualization of cell migration through the conduit into the left chamber [32] (**Figure 4A**, supplementary **Movie S1**). Migrating cells were morphologically distinct compared to non-migrating ones (**Figure 4Bi,ii;** supplementary **Movies S2, S3**).

**Figure 4.**
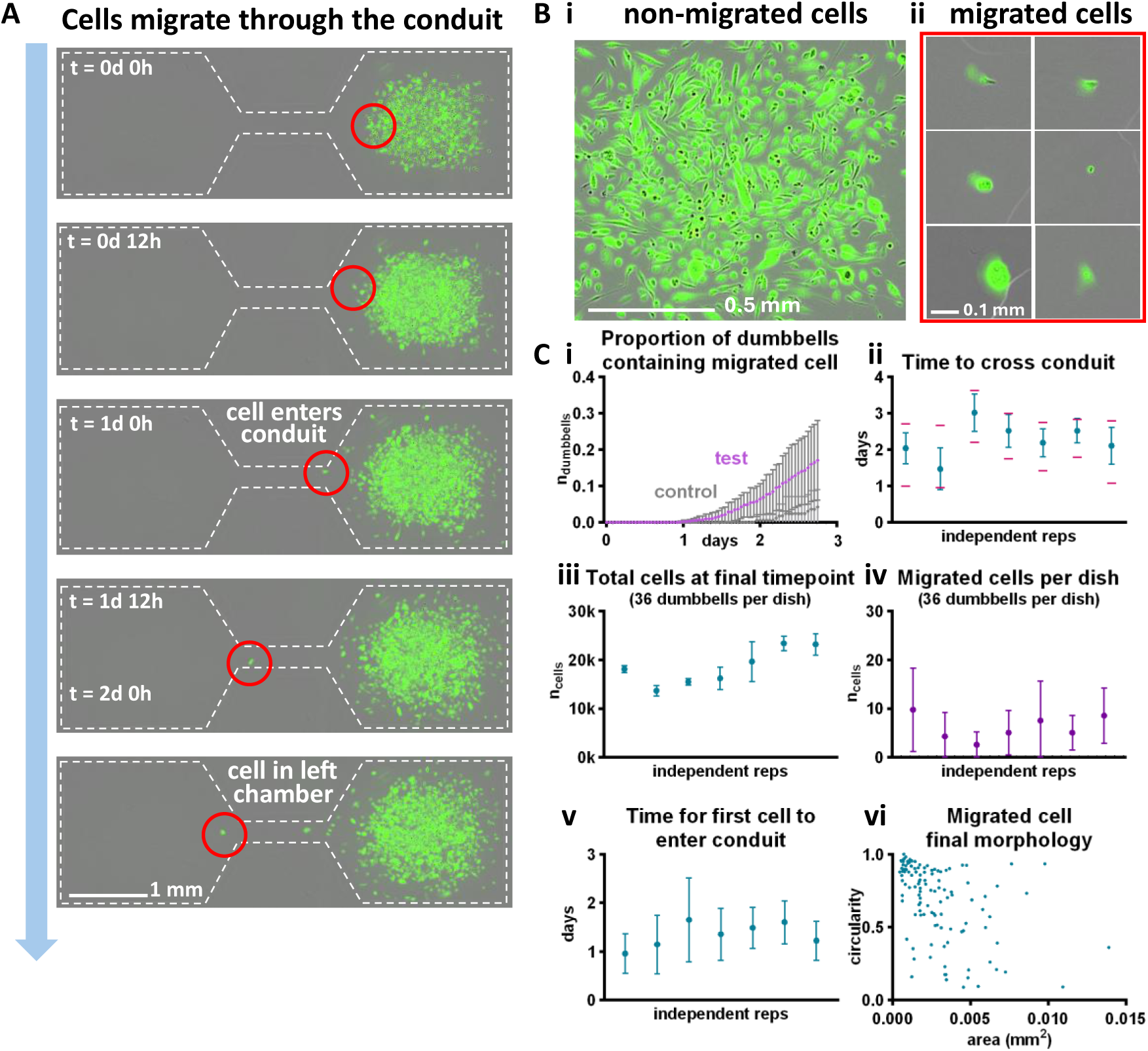
Single-cell migration through dumbbells. **(A)** PC3-eGFP cells are initially confined to the right chamber. Excellent circuit optics allows migration of single cells to be followed. **(B)** Representative fluorescence images showing cell morphology in **(i)** the non-migrated cell population and **(ii)** migrated cells at the final timepoint. **(C)** Some key migration parameters across 7 independent replicates. **(i)** Proportion of dumbbells on a dish containing a migrated cell over time. **(ii)** Times taken for a cell to traverse the conduit, with max/min time taken shown as pink lines. **(iii)** Total number of cells across all 36 active dumbbells on a dish at the final timepoint. **(iv)** The average number of migrated cells on a dish for each replicate. **(v)** Average time taken for the first cell to enter the conduit. **(vi)** Morphologies of migrated cells at the final timepoint. Error bars in each sub-panel indicate standard deviations. Each experimental replicate comprised 120 dumbbells split evenly across 4 dishes.

Analysis of time-lapse image stacks of migrating cells revealed key migration parameters. Data are presented across 7 independent replicates (experiments performed on different days using the same cell line), with each experiment comprising 120 test dumbbells and 24 control dumbbells (no FBS), distributed equally across four dishes (36 dumbbells per dish, the maximum number visible within the field of view of the imaging system).

In the test condition, cells required at least 24 h to traverse the conduit, with the proportion of dumbbells containing migrated cells increasing steadily over 3 days (**Figure 4Ci**). Migration was significantly greater in the test condition than in the control (p < 0.01, Welch’s two-sided t-test comparing area under the curve), indicating that diffusion of FBS through the conduit effectively promotes cell migration. The average time required for a cell to traverse the conduit was typically 2-3 days (**Figure 4Cii**). Comparing the total number of cells in all 36 dumbbells on any dish at the final timepoint **(Figure 4Ciii**) with the number of migrated cells per dish (**Figure 4Civ**) reveals that migrated cells represent **⁓** 0.03% of the total cell population, averaged across all 7 replicates.

There was substantial variability in the time required for the first cell to enter the conduit, with differences of days observed between dumbbells on the same dish (**Figure 4Cv**). In some cases, cells entered the conduit in <5 h – shorter than the time required to establish a stable FBS gradient (**⁓** 6 h) – suggesting that early entry reflects exploratory behaviour rather than chemotaxis. Approximately 30% of dumbbells contained cells that followed the first cell through the conduit; followers were deliberately excluded from analysis to focus on the most rapidly-migrating ones.

Migrated cells were morphologically heterogeneous (**Figure 4Cvi**) with areas ranging from 0.00043 mm^2^ (smallest) to 0.0139 mm^2^ (largest). Of the 119 cells measured, 48 had an area below 0.004 mm^2^ and a circularity above 0.8, indicating a substantial fraction of small, round cells at the final timepoint relative to a mid-range value of 0.0067 mm^2^. The remaining cells displayed diverse morphologies, including larger, elongated, and irregularly-shaped phenotypes.

Conduit height depends on Laplace pressure, assumed to be equal to that of the left and right chambers at equilibrium. Low-volume conditions (<200 nL chamber volume) enabled the formation of very shallow conduits (see supplementary **Figure S2** for detailed theory). Assuming equilibrated chamber volumes of **⁓** 150 nL, equating pressures yields an estimated conduit height of ⁓ 1.3 µm, smaller than the height of adherent PC3 cells (⁓ 3 – 5 µm [34]). Similarly shallow heights have been reported in comparable open microfluidic migration assays employing silicon oil-medium interfaces, where interfacial deformation was observed during cell transit [14]. A similar interaction is likely to occur here; interfacial deformation may impose mechanical constraints on migrating cells.

### Reconfiguring dumbbells to isolate and pick cells of interest

A key advantage of fluid walls is the ability to dynamically reconfigure geometries during an experiment [18], [25]. We applied this capability to demonstrate how the most migratory single cells can be isolated from non-migrated ones. When a single migrated cell was identified in the left chamber, the same FC40 jet used for dumbbell fabrication was redeployed to construct new fluid walls across the conduit entrance and exit, thereby isolating each chamber. This required a higher velocity FC40 jet than used to initially form the fluid walls, to ‘cut’ through the conduit, which has (at equilibrium) a cross-sectional area of **⁓** 170 μm^2^. Consequently, the jet flow rate was increased from 480 μL/min to 900 μL/min to ensure sufficient FC40 could be jetted vertically through medium in the conduit onto the bottom of the dish to form the fluid wall. Following isolation of the target cell from all the others, trypsin was used to detach the cell from the polystyrene surface, and – by withdrawing volume – the cell was picked and transferred to a microtiter well (**Figure 5A**). An additional step was required to obtain single non-migrated cells. Non-migrated cells were isolated, detached, and pooled, and the resulting cell suspension distributed across a GRID of fluid-walled micro-chambers [35] (**Figure 5B**). If a single cell was identified within a chamber, it was picked using an optimized method (supplementary **Figure S8**).

**Figure 5.**
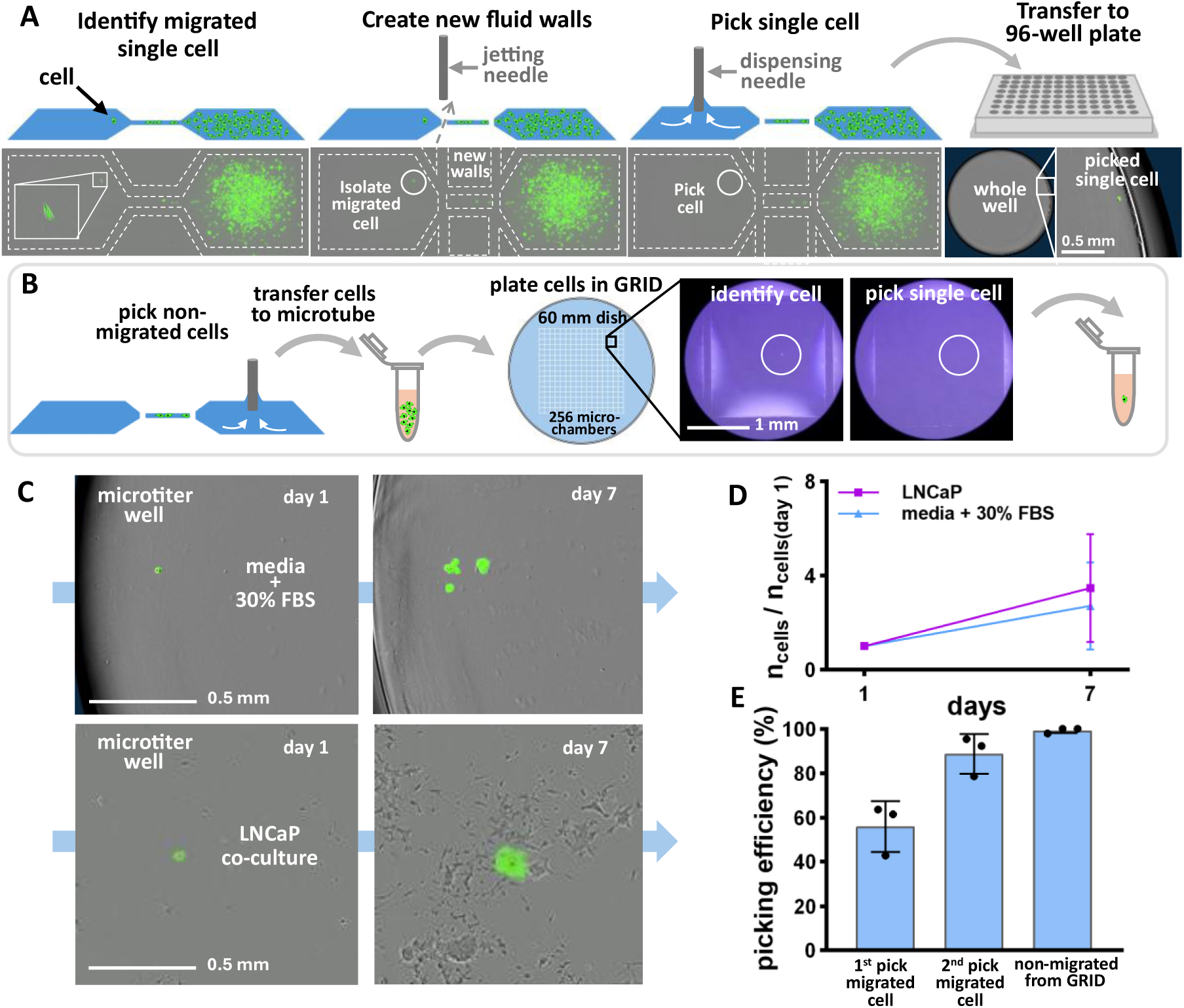
Reconfigure circuit to isolate and pick single cell of interest. **(A)** Workflow for isolation and picking of migrated cell (PC3-eGFP) from dumbbells. Fluorescence-phase images of dumbbells shown below. First, a single migrated cell is identified. Then, a jet of FC40 is used to create new fluid walls across the conduit, isolating the cell from the non-migrated population. The cell is then picked from the chamber and transferred to a microtiter well. **(B)** Workflow for picking many non-migrated cells, followed by clonal isolation. Non-migrated cells are picked and collected in a microtube, then the suspension is plated across a GRID of fluid-walled chambers, single cells identified and picked for downstream analysis. **(C)** Single migrated cells are picked and transferred to microwells containing medium + 30% FBS, or LNCaP co-cultures. **(D)** Cell proliferation in different environments, quantified as fold increase in cell number at d 7 relative to counts one day after picking (± standard deviation, 20 cells transferred to LNCaP co-culture and 25 to medium + 30% FBS). **(E)** Comparison of single-cell picking efficiency. Cells picked by two sequential rounds of picking showed high efficiency, comparable to the optimized GRID-based method (± standard deviation, n = 3).

Proliferation of picked single migrated cells was assessed under two conditions: medium + 30% FBS and co-culture with LNCaP cells. Migrated cells grew to yield clones (**Figure 5C**), with an average cloning efficiency of 12% in the high-serum condition – lower than that achieved by other single-cell handling systems [36] and likely due to the relative difficulty in lifting cells from the FNC-mix-coated surface, requiring trypsinisation for > 15 mins and often two consecutive rounds of picking. Quantification of fold-change in cell number at day 7 revealed a numerically higher mean normalised cell recovery in co-culture, although substantial variability was observed among technical replicates (**Figure 5D**), with 20 migrated cells transferred to co-culture and 25 to medium + 30% FBS. Picking efficiency was also evaluated (**Figure 5E**). Approximately half of migrated cells (3 replicates) were picked on the first attempt, and re-picking missed cells raised recoveries comparable to those found in our optimized GRID-based workflow (**Figure S8**).

### AI segmentation and tracking of single cells

We exploited the excellent optics provided by the transparent fluid walls to train a custom model for automated segmentation and tracking of individual cells using Cellpose-SAM. Training on 100 manually-segmented images was sufficient to reach ⁓97% agreement with manual counting (**Figure S9**). Hourly live-cell imaging enabled high-resolution single-cell tracking and extraction of dynamic morphological features during migration (**Figure 6A,B**). Using this trained model in combination with ImageJ, cells were tracked across 58 representative dumbbells, each containing a single migratory cell (supplementary **Movie S4, S5**). If multiple cells migrated through the conduit, analysis was restricted to the trajectory of the first cell to complete transit. Analysis of cell trajectories using polar histograms (rose plots) revealed a clear directional bias of migrated cells towards and through the conduit (**Figure 6Ci**), whereas non-migrating ones showed no such bias (**Figure 6Cii**).

**Figure 6.**
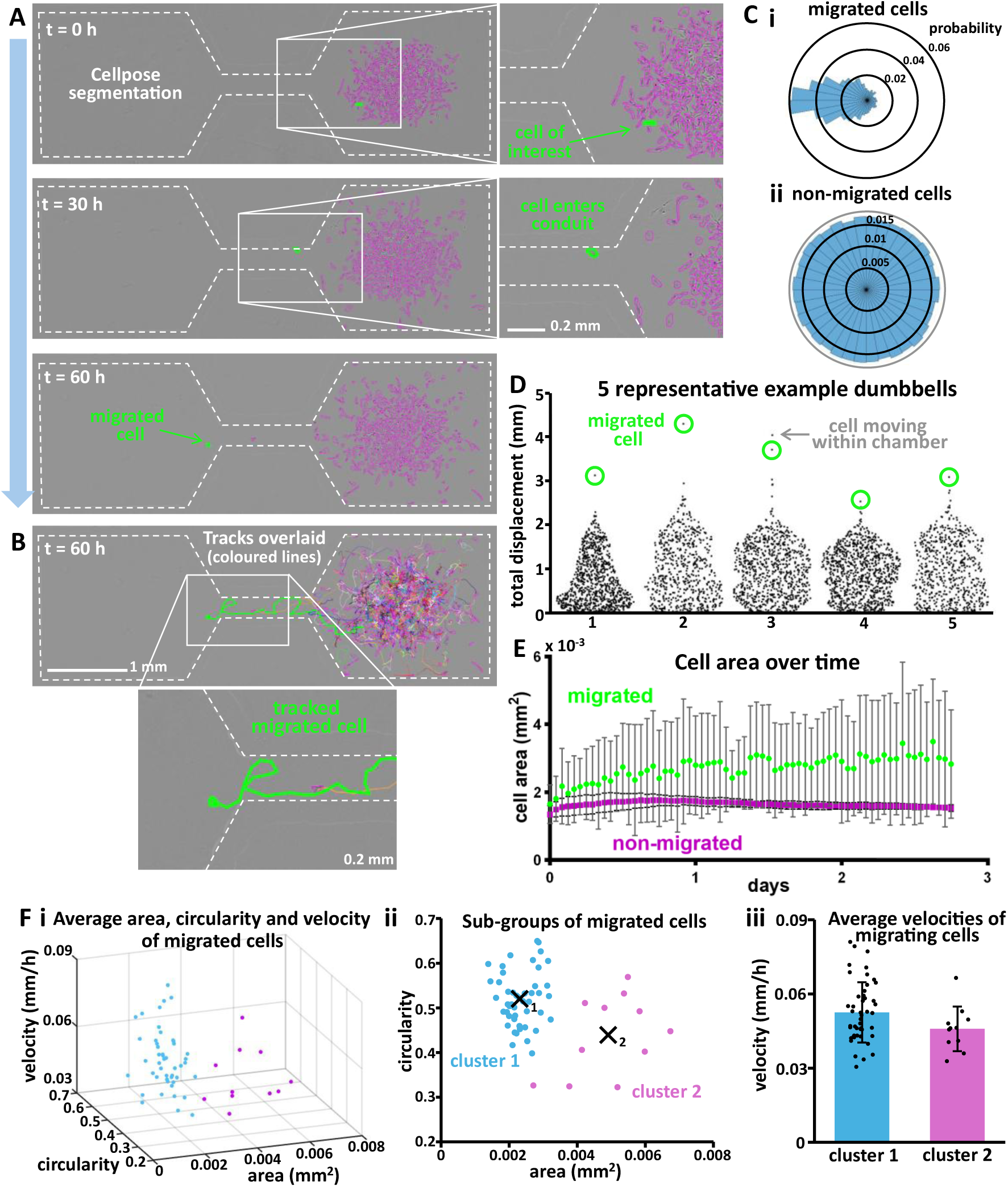
Segmentation and tracking of single cells. **(A)** A custom Cellpose-SAM model, trained on representative dumbbell images, was used to segment individual cells in time-lapse stacks. Hourly imaging enabled high-resolution cell tracking. A green line is drawn around the cell of interest, and purple around all others. **(B)** Final frame from A with tracks overlaid. The migrated cell is highlighted in green, with multicoloured lines showing tracks of non-migrated cells. **(C)** Migrated cells showed a strong bias towards the left, through the conduit **(i)**, whereas non-migrated cells had no directional preference **(ii).** Data shown for 58 migratory cells and ⁓3 000 non-migrated. **(D)** Comparison of track lengths revealed that migrated cells typically had the greatest total displacement; however, in some cases, highly-motile cells failed to traverse the conduit. Data from 5 representative dumbbells are shown. **(E)** Integration of segmentation with tracking allowed quantification of changes in cell area in migrated versus non-migrated populations over time (± standard deviation, 21 migrated cells and ⁓10 000 non-migrated). **(F)** Comparison of **(i)** average area, circularity, and velocity among migrated cells. Cluster analysis based on **(ii)** cell morphology reveals distinct subpopulations, which have **(iii)** different motility characteristics (± standard deviation, 47 cells in cluster 1 and 11 in cluster 2, p = 0.054, Welch’s two-sample t-test).

We next compared the total displacement of each tracked cell in 5 representative examples. In most cases, the cell that successfully migrated through the conduit also exhibited the greatest overall displacement within the circuit. However, we also observed instances in which a cell exhibited high motility within the chamber but did not traverse the conduit (**Figure 6D**).

Combining segmentation with tracking enabled assessment of morphological dynamics during migration. For example, analysis of cell area revealed that both migrated and non-migrated populations began with similar sizes (average cell area of 0.0017 mm^2^ and 0.0013 mm^2^ respectively), but migratory cells exhibited a greater increase in area over time compared with the average non-migrated cell (final average cell areas of 0.0028 mm^2^ and 0.0015 mm^2^ respectively, with maximum average values of 0.0035 mm^2^ and 0.0018 mm^2^ for each group, **Figure 6E**).

To relate morphology to migratory behaviour of 58 migratory cells, we compared average cell area, circularity, and velocity (**Figure 6Fi**). Visualisation of area against circularity revealed that migrated cells segregated into distinct morphological subpopulations. K-means clustering identified two groups: a population of smaller, more circular cells and a population of larger, less-circular ones (**Figure 6Fii**). The first group had an average cell area of 0.0023 mm^2^ and circularity of 0.52, whereas the second had an average area of 0.0049 mm^2^ and circularity of 0.44. Analysis of motility revealed that the smaller, circular cells migrated at higher average velocities (0.053 mm/h) compared with the larger, less-circular counterparts (0.046 mm/h, **Figure 6Fiii**, supplementary **Movie S6**). See supplementary **Figure S9** for further information on implementing Cellpose and ImageJ for automated segmentation and tracking of single cells.

## Discussion

Cancer metastasis remains a major clinical challenge. We present an accessible fluid-walled microfluidic platform for biologists to create high throughput migration assays on-demand, overcoming many of the fabrication barriers of conventional microfluidics. Jet-printing enabled rapid and reproducible dumbbell fabrication in seconds, with all regions fully accessible through oil-medium interfaces (**Figure 1**). Dumbbell stability requires protein adsorption to the media-FC40-polystyrene pinning line [31], and 0.5% FBS concentration allowed reliable fabrication (**Figure S1**). Modulating circuit volumes enabled control of flow direction (**Figure 2**) and spatial confinement of cells and chemo-attractant to distinct dumbbell chambers (**Figure 3; Figure S4**). Over 3 days, some cells migrated through the conduit towards serum (**Figure 4**). This is consistent with findings from conventional solid-walled microfluidic migration assays, which have shown that within heterogenous cancer cell populations, only a subset of cells exhibit migratory behaviour through microchannels [11], [12]. Within dumbbells, only a small minority of cells migrated, with a dish containing 36 dumbbells yielding on average only 3 migratory cells; this aligns with the current thinking that only a small fraction of cells within a primary tumour are likely metastatic [37]. Migratory cells were morphologically heterogeneous, ranging from small and round to larger cells extending filaments resembling lamellipodia forming in the direction of movement (**Figure 4B**). Media + 20% FBS promoted migration (**Figure S6**), and smaller volumes limited proliferation, ensuring that cells in the conduit were migrating rather than expanding (**Figure S7**). Cells often explored the chamber before entering the conduit, however in some cases cells could be observed immediately moving towards and through the conduit (**Movie S1**). As some cells migrated in serum-free controls, serum enhances, rather than initiates, migration in an inherently exploratory subpopulation (**Figure 4Ci**).

Conduit height depends on Laplace pressure, with low chamber volumes (and hence pressure) enabling very shallow conduits (e.g. ⁓ 1.3 µm, with 150 nL in each chamber), smaller than the height of many adherent cells. Many PDMS-based microfluidic migration assays employ similar channel heights (⁓ 3 – 5 µm) to prevent cells in suspension from entering microchannels, or mimic aspects of metastasis where cells navigate through restricted microenvironments [11], [12], [13], [38], [39]. Even shallower channel heights (< 1 µm) were reported in a previous study employing oil-medium interfaces [14]. In these systems, interfacial deformation during cell transit imposed compressive forces that elicited bleb-based migratory behaviour, and more closely recapitulated migration through confined tissue environments than solid-walled systems. A similar interaction is likely to occur in the present fluid-walled assay, where interfacial deformation may impose mechanical constraints on migrating cells, or influence processes associated with the epithelial-mesenchymal transition (EMT) and hence cell mobility [40]. Accordingly, future work could include analysis of EMT markers in migratory versus non-migratory cells to further elucidate these effects.

Single migratory cells were isolated, picked, and yielded clones (**Figure 5**), however cloning efficiency was low compared to other automated single-cell handling systems – further optimisation is required to improve clonal recovery. We also show a method for obtaining single non-migrated cells, demonstrating a comparative framework (**Figure 5B**). The ability to isolate and retrieve single migratory cells represents a key advantage of fluid walls over conventional microfluidic platforms, where migratory or non-migratory cells are typically pooled for downstream analysis [11], [12], [13]. While such approaches enable comparisons between broad groups, they may overlook heterogeneity within migratory subpopulations.

The transparency of fluid walls enabled the collection of images throughout the workflow, and the use of automated segmentation and tracking routines allowed comparison between both the morphologies and motilities of migratory and non-migratory cells (**Figure 6**). While the Cellpose/ImageJ tracking worked well, it was computationally expensive, taking ⁓5 minutes per dumbbell on GPUs and ⁓40% of tracked migratory cells required manual correction. The migrating cells grew more rapidly over time than non-migrating ones (**Figure 6E**) likely due to entering a serum-rich environment. K-means clustering was used to identify two migratory groups: smaller, rounder cells and larger, less circular cells (**Figure 6F**). On average, smaller cells migrated faster than larger ones, although the difference between each group did not reach statistical significance, likely due to the limited sample size (47 and 11 cells respectively, p = 0.054, Welch’s two-sample t-test). This is likely due to differences in traction energy associated with attachment area [41]. Furthermore, we cannot directly infer that faster-moving cells possess greater metastatic potential as cells may adopt different migratory behaviours in more complex/3D environments. Nevertheless, prior studies have reported correlations between increased motility and metastatic fitness [12], [13], [42].

While our platform has many advantages, some limitations should be noted. Experimental conditions were optimised for PC3 cells; different cell types would require re-optimisation to ensure sufficient yields. Furthermore, diffusion gradients exist only within the conduit, making it challenging to infer precise chemotactic responses, although alternative fluid-walled designs exist for such applications [19], [43]. In summary, this work establishes fluid-walled microfluidics as an accessible platform for studying single migratory cancer cells, opening new avenues for dissecting the functional heterogeneity underlying metastasis. Metastasis is thought to be driven by rare and phenotypically heterogenous tumour-cell populations, with increasing evidence for single-cell biology to predict metastatic potential [44], [45], [46]. As such, effective approaches to track and interrogate such cells provides a powerful opportunity to further our understanding of the biology underlying cancer metastasis.

## Materials and Methods

### Cell culture

Cells were grown routinely at 37°C, 5% CO_2_, 20% O_2_ and live cells counted using a Bio-Rad (Bio-Rad Laboratories, Hercules, CA, USA) TC20 automated cell counter with Trypan Blue dye exclusion unless stated otherwise.

PC3-EGFP cells [21] and LNCaP cells [47] were kind gifts from Yolanda Calle (Kings College London) and Dr Richard Bryant (University of Oxford, Oxford, UK) respectively. Human prostate cancer cells were cultured in 20 mL RPMI-1640 medium (Sigma-Aldrich, St. Louis, MO, USA) + 10% FBS (Gibco, ThermoFisher Scientific, Waltham, MA, USA) + 1% MEM nonessential amino acids (Gibco) + 1% penicillin-streptomycin (Gibco) + 1% L-Glutamine (Gibco) + 1% sodium pyruvate (Gibco) + 1% MEM vitamin solution (Gibco) in 75 cm^2^ U-shaped canted neck flasks (Corning). We refer to this media composition as ‘complete’ media (without FBS). Cells were passaged or prepared for experiments by first pre-warming media, PBS (Sigma-Aldrich) and trypsin (TrypLE Express, Gibco) in a 37°C water bath. Media was removed, cells rinsed (PBS, 3 mL) and detached (trypsin, 3 mL). Cells were incubated (3 mins), 4 mL media added, suspension transferred to 15 mL centrifuge tube (300g; 5 min) and supernatant removed. Cells were resuspended in 5 mL media, counted and diluted to the required concentration.

Human embryonic kidney (HEK 293) cells [48] were cultured in 5 mL DMEM (Sigma-Aldrich) + 10% FBS + 1% penicillin-streptomycin in 25 cm^2^ polystyrene canted neck flasks (Corning). Cells were passaged or prepared for experiments as above with the following modifications: rinsed with 1 mL PBS, detached (trypsin, 1 mL), 4 mL media added then spun down (500 rpm; 3 min). Cells resuspended in PBS and counted using a haemocytometer.

### Fluid printer

The fluid printer (iotaSciences Ltd) comprised a three-axis traverse and two integrated syringe pumps used to automate fluid deposition via two needles (one per syringe). The first syringe contained FC40 and connected to a needle (70 μm internal diameter) spatially controlled via the traverse to jet-print the microfluidic circuit; the ‘jetting needle’. The second was filled with 70% ethanol solution (to maintain sterility) and connected to the ‘dispensing needle’ (255 μm internal diameter) used in the handling of media, cells and reagents. Needles were connected to respective syringe pumps via Teflon tubing. The central platform within the printer held a single 60 mm petri dish and up to eight 1.5 mL Eppendorf tubes. A custom platform was attached to the front of the printer to accommodate microtiter well plates, allowing automated transfer of cells from petri dishes.

### Fabrication and preparation of dumbbells with fluid walls

Dumbbells with fluid walls were jet-printed following the workflow outlined in the experimental section of this paper, creating arrays of 80 dumbbells (5 columns, 16 rows, for detailed geometry, see **Figure S2**). The jetting process swept aside the underlying media such that each dumbbell contained approximately 150 nL (estimated by comparing weights of empty dishes and those covered by a thin film of media). Then, 600 nL PBS was added to the left and right chambers of the inner 36 dumbbells (the field of view of the live-cell imaging system) and 800 nL subsequently removed – thereby also withdrawing a small volume of FC40. This minimized variability in dumbbell volume post-jetting and washed away residual serum (effectiveness of wash shown in **Figure S3**). Dumbbell chambers were then filled with 100 nL human FNC coating mix (AthenaES) to encourage initial cell attachment.

### Enabling spatial confinement of cells and chemo-attractant

First, 400 nL complete serum-free media was added to the left chamber (500 nL total volume). Then, 100 nL suspension (⁓ 500 cells in serum-free complete media) added to the right chamber (200 nL total volume). Differences in chamber volumes ensured flow from the left chamber to the right and hence cell confinement (**Figure S4**). Dishes were incubated for ∼ 3 h to allow cells to attach. Before adding chemo-attractant, 400 nL was removed from the right (cell) chamber and 600 nL removed from the left, such that flow was always in the direction of the cell chamber. Then, 200 nL complete serum-free media was added to the cell chamber, and 100 nL complete media + 20% FBS was added to the left. Cell suspensions were aspirated manually before collection by the dispensing system, mitigating sedimentation within the storage tube.

### Live cell imaging during migration

Following addition of chemo-attractant, cells were imaged hourly for ⁓ 3 days using the Incucyte S3 live-cell analysis system (Sartorius, Göttingen, Germany), acquiring phase-contrast and fluorescence (green) scans of the 36 inner dumbbells. Dishes were positioned using a custom holder (iotaSciences) and were typically loaded four at a time. Image data were organised using MATLAB (2023a, MathWorks, Natick, MA, USA) and time-lapse sequences of cell migration (phase-contrast and fluorescence) were created using ImageJ. To correct for positional drift, each cropped stack was stabilised using the template-matching plugin in ImageJ. Cell images in **Figures 3-5** are combined phase and fluorescent, **Figure 6** is phase only.

### Optimization of cell number, chamber volume, and chemo-attractant concentration

To optimize seeding concentrations used, dumbbells were fabricated as usual, and each dish divided into 3 columns of 12 dumbbells. Cell suspension (100 nL) was seeded into the right chamber at ⁓125, 250, or 500 cells. For each condition, cells were scored if they entered the conduit or progressed over halfway. The highest concentration (⁓500 cells per dumbbell) yielded the most cells progressing over halfway through the conduit (**Figure S5**).

To assess the effect of serum concentration and chamber volume on migration, dumbbells were fabricated and seeded with ⁓500 in the right chamber. After cell attachment, medium containing varying FBS concentrations (0% – 100%, 20% increments) was added to the left chamber. Dumbbells were arranged in 6 groups of 6 per dish, with experiments performed across 4 dishes: 2 with equilibrated chamber volumes ⁓150 nL, and 2 ⁓450 nL. Dishes were imaged hourly using the Incucyte, and cells scored if they entered the conduit or progressed over halfway. Total cell number within a circuit was counted using ImageJ (**Figure S7**).

### Automated cell segmentation and tracking

Hourly phase-contrast images acquired using the IncuCyte were compiled into time-lapse stacks using MATLAB/ImageJ and stabilized. Stacks were cropped to single dumbbells containing cells migrated through the conduit. From each stack, the 20^th^, 30^th^ and 40^th^ frames were extracted as representative samples for model training. Training images were sequentially loaded into the Cellpose graphical user interface (GUI), where the Cellpose segment anything model (SAM) provided initial segmentations. Segmentation errors were corrected manually for a human-in-the-loop workflow. In total, 150 images were annotated and custom models trained on an NVIDIA Quadro RTX 400 GPU (NVIDIA, Santa Clara, CA, USA). Subsequently, full time-lapse stacks were processed in ImageJ using the TrackMate plugin with the Cellpose detector module. Cell trajectories were generated using the Linear Assignment Problem (LAP) tracker. For the primary cell of interest in each stack, any tracking errors were manually corrected. The results datasets (including X-Y coordinates, cell area, perimeter, and circularity at each timepoint) were exported for further analysis.

### Picking single cells post-migration

After 3 days, dishes were removed from the incubator and transferred to the fluid printer. To isolate the most migratory cells from the non-migrated population, new fluid walls were jet-printed intersecting both ends of the conduit (‘cutting’), physically separating the two populations. To ensure consistent cutting, a higher jet rate of 900 μL/min was used. Then, dishes were imaged to confirm successful isolation and identify suitable candidates for retrieval. For each target chamber, 400 nL PBS was added and 600 nL subsequently removed to minimize chamber volume. Cells were detached using 400 nL trypsin per chamber, 20 mins incubation. The entire chamber volume was withdrawn, and the retrieved cell transferred to a 96-well microtiter plate with wells pre-filled with 100 μL media. Chambers were re-imaged after picking to confirm the removal of the target cell. If the cell remained, an additional 200 nL trypsin was added and the picking process repeated.

The same procedure was then applied to the non-migrated cell population, with the following modifications. Non-migrated cells (from dumbbells containing a migrated one) were picked into the first row of a 96-well plate, each well pre-filled with 100 μL media. The suspension was then pooled into a 1.5 mL tube. In parallel, a microfluidic GRID comprising 256 microfluidic chambers was created [35]. A 60 mm dish was coated with a thin layer of PBS + 5 mg/mL BSA, overlaid with 2 mL FC40, and the fluid printer used to jet-print the GRID. Chambers were pre-filled with 100 nL PBS. The pooled cell suspension was then dispensed into the GRID, 100 nL per chamber. Visual inspection via microscope was used to confirm the presence of a single cell in a chamber. Chambers containing a single cell were subsequently aspirated and the cell transferred to a microtiter well (pre-filled with medium).

### Optimization of Single Cell Picking from GRID chambers

Arrays of 256 square chambers were jet-printed in 60 mm tissue culture-treated polystyrene dishes (Corning). To quantify picking efficiency, ⁓ 100 cells were seeded into each chamber. Cells were counted before and after picking, and efficiency defined as percentage of cells removed. The underlying medium used in GRID fabrication was varied, including 0.05%, 0.5%, and 5% BSA in PBS (to test whether BSA, known to inhibit cell adhesion, improved picking [49]), as well as standard culture medium supplemented with 10% FBS.

To ensure cells remained centrally located within chambers, we tested different ratios of chamber pre-fill volume to suspension added. Chambers were pre-filled with 50 or 100 nL PBS, after which cell suspension was added (200 nL total chamber volume). Cells were imaged (Olympus CKX53, Olympus, Tokyo, Japan) and positions measured relative to chamber centre using ImageJ. Optimization experiments were first performed with HEK and LNCaP cells before PC3. Increasing BSA concentration and pre-filling chambers with PBS improved picking efficiency, the optimized method required chamber fabrication with PBS + 5 mg/mL BSA, pre-filling with 100 nL PBS, and adding 100 nL cell suspension (**Figure S8**).

### Visualising confinement and diffusion of chemo-attractant

Dishes and dumbbells were prepared as usual. After the final wash step, 200 nL PBS was added to the chamber that would normally contain cells. Subsequently, 100 nL complete media + 10 mg/mL BSA was added to the opposing chamber. The BSA solution comprised a mixture of fluorescently labelled and unlabelled protein at a 1:10 ratio (BSA AlexaFluor 488 conjugate (Invitrogen, ThermoFisher Scientific, Waltham, MA, USA); BSA Fraction V, Sigma-Aldrich), corresponding to an effective FBS concentration of 20%.

Dishes were imaged every 2 h for 2 d using the Incucyte (phase and GFP fluorescence images), and time-lapse image stacks created as before. Time required for fluorescent albumin to enter the conduit was determined by drawing a line profile across the junction between the chamber and conduit and a second line 100 μm into the conduit in ImageJ, and monitoring pixel intensity to detect GFP signal. Estimated diffusion rates were calculated in Excel (version 2404, Microsoft, Redmond, WA, USA).

### Visualising effect of PBS wash in dumbbells

To assess the efficiency of washing in removing residual serum, 60 mm dishes were coated with 1 mL complete media + 0.25 mg/mL fluorescent BSA. The coating solution was withdrawn and overlaid with 2 mL FC40. Dumbbells were then jetted as normal and dishes imaged using the Incucyte. For the central 36 dumbbells, 600 nL PBS was added to the central and right columns (24 dumbbells) followed by the withdrawal of 800 nL to complete the first wash. Dishes were re-imaged, after which the right column (12 dumbbells) underwent an additional wash before imaging a final time. Fluorescent intensity within each dumbbell was quantified using ImageJ.

### Estimation of circuit local volumes and flow rates

Dumbbells matching the geometry used in experimental assays were fabricated in large uni-well dishes (Nunc^TM^, ThermoFisher Scientific, Waltham, MA, USA) with flat walls, improving side-on imaging quality compared to curved walls of Petri dishes. Dishes were coated with 4 mL DMEM + 0.5 % FBS; medium was withdrawn to leave a thin film, and ⁓ 20 mL FC40 overlaid. Since the normal fluid printer cannot accommodate larger dishes, a custom three-axis traverse system (iotaSciences) was used. The jetting needle was connected to a syringe pump (kd Scientific, Holliston, MA, USA) driving a 2 mL syringe (Hamilton, Reno, NV, USA) filled with FC40. A separate syringe pump (PhD Ultra, Harvard Apparatus, Holliston, MA, USA) controlled a 50 μL syringe (Hamilton) connected to a dispensing needle. After jetting, 100 nL serum-free media was dispensed into each chamber, and the dish imaged from side-on (FTA1000 B Class, First Ten Angstroms, Newark, CA, USA). Next, 400 nL of media was added to the left chamber and 100 nL to the right, with images acquired every 5 mins for 6 hours. Chamber heights were determined using FTA32 software, with the dispensing needle external diameter (0.51 mm) as a scale reference. Chamber volumes and corresponding pressures were estimated from measured heights. Flow rate was estimated from the curve fit to change in average height of the taller drop over time (curve fit to the average height was ℎ(*t*) = 0.0119*e*^−1.1041*t*^ + 0.2346*e*^−0.016*t*^ where ℎ is drop height in mm and *t* time in hours).

### Statistical analysis

All calculations, including estimation of local pressures within dumbbells, 3D geometry and diffusion rates, and statistical analyses were performed using Excel (Microsoft) or MATLAB (MathWorks). K-means clustering was performed on z-score-standardized area and circularity data to identify morphological subpopulations (k = 2). Differences in mean migration velocity between resulting clusters were assessed using a two-sample Welch’s t-test. A p-value below 0.05 was considered significant.

## Acknowledgements

This work was supported by the Medical Research Council (grant number MR/N013468/1) and Rosetrees and The John Black Charitable Foundation (grant numbers PGS21/10073 and CF-2024-M-2\114). The authors also thank iotaSciences (iota Sciences Ltd, Begbroke Science Park, Oxfordshire, OX5 1PF) for providing equipment, reagents and financial support toward the project.

## Notes

**Competing interest statement:** P. R. C. and E. J. W. co-founded, and hold equity in, IotaSciences Ltd who provide support to J.A.E.M and equipment for the study.

### Competing Interest Statement

P. R. C. and E. J. W. co-founded, and hold equity in, IotaSciences Ltd who provide support to J.A.E.M and equipment for the study.

