## Supplementary Information for "Identification and isolation of migratory subpopulations of cancer cells using microfluidics with fluid walls"

### Serum content required for fabrication of fluid walls

BSA is an excellent stabilizer of fluid walls, and RPMI medium plus FBS (which contains BSA) is used as a chemo-attractant. Consequently, FBS-free medium is applied as the control, and it is then important that this contains too little FBS to act as an attractant. Therefore, when jetting, we employed RPMI medium plus the lowest FBS concentration sufficient to stabilize footprints during subsequent volume additions (e.g., during ‘washing’).

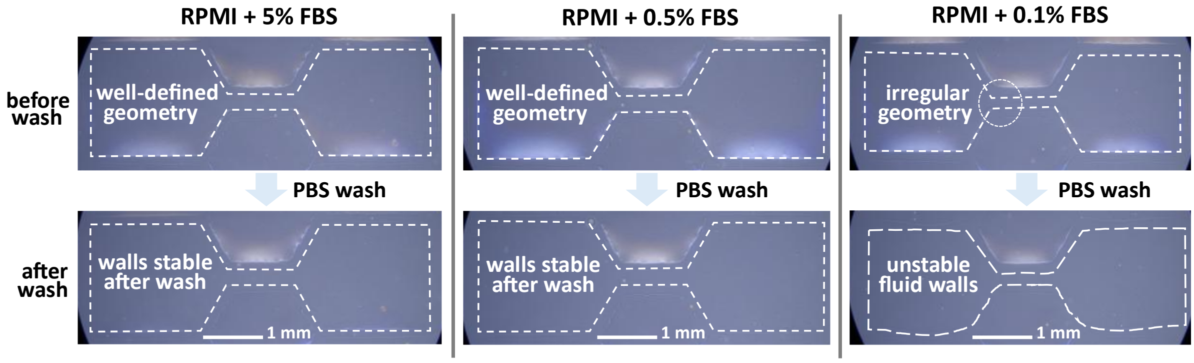


**Figure S1. Serum content required for fabrication of fluid walls.** Dashed white lines mark microenvironment footprints. Footprints are formed correctly after jetting through RPMI plus three dilutions of FBS (top). After washing each chamber with 600 nl PBS, footprints made with RPMI + 0.1% FBS, prove to be unstable (see curved pinning lines at bottom right and notably narrower conduit at top right, highlighted with circle).

### Microfluidic dumbbells with fluid walls: pressure dynamics, 2- and 3-D geometries

Fluid-walled environments are bounded by liquid-liquid interfaces. While their 2-D geometry is stable, 3-D geometry morphs when volumes are added or subtracted. Our dumbbell chambers are sufficiently small that interfacial forces dominate and govern their 3-D curvature. To estimate the radius of curvature of chambers, we approximate the Laplace pressure by equating it to that of a circular droplet with an equivalent footprint area. The equivalent footprint radius $a_{eq}$ is defined as:

$$a_{eq} = \sqrt{\frac{chamber\ area}{\pi}} \tag{S1}$$

This approximation allows use of spherical-cap geometry in relating chamber height to volume $V$:

$$V = \frac{1}{6}\pi h\left(3a_{eq}^2 + h^2\right) \tag{S2}$$

and Laplace pressure:

$$P_L = \frac{2\gamma}{R} = \frac{4\gamma h}{h^2 + a_{eq}^2} \tag{S3}$$

Total pressure in the chamber is then given by summing Laplace and hydrostatic effects due to media and FC40:

$$P = \frac{4\gamma h}{h^2 + a_{eq}^2} + \rho_{media} g h + \rho_{FC40} g h_{FC40} \tag{S4}$$

The height of FC40 is calculated using a dish surface area of 21.5 $cm^2$ and 3 ml overlay. At equilibrium, the Laplace pressure in the chamber is equal to the pressure in the conduit where $P_{conduit} = \gamma/R_{conduit}$. Furthermore, we assume the height of the conduit to be much less than the height of the drop or conduit width ($h_c \ll a$ and $h_c \ll h$). This gives the relationship between chamber and conduit height:

$$h_c = \frac{2a_c^2 h}{h^2 + a_{eq}^2} - \frac{\Delta\rho g h a_c^2}{2\gamma} \tag{S5}$$

This can be used to calculate the cross-sectional area of the conduit, $A_c$:

$$A_c = \frac{(a_c^2 + h_c^2)^2}{4h_c^2}\cos^{-1}\left(\frac{a_c^2 - h_c^2}{a_c^2 + h_c^2}\right) - \frac{a_c}{2h_c}(a_c^2 - h_c^2) \tag{S6}$$

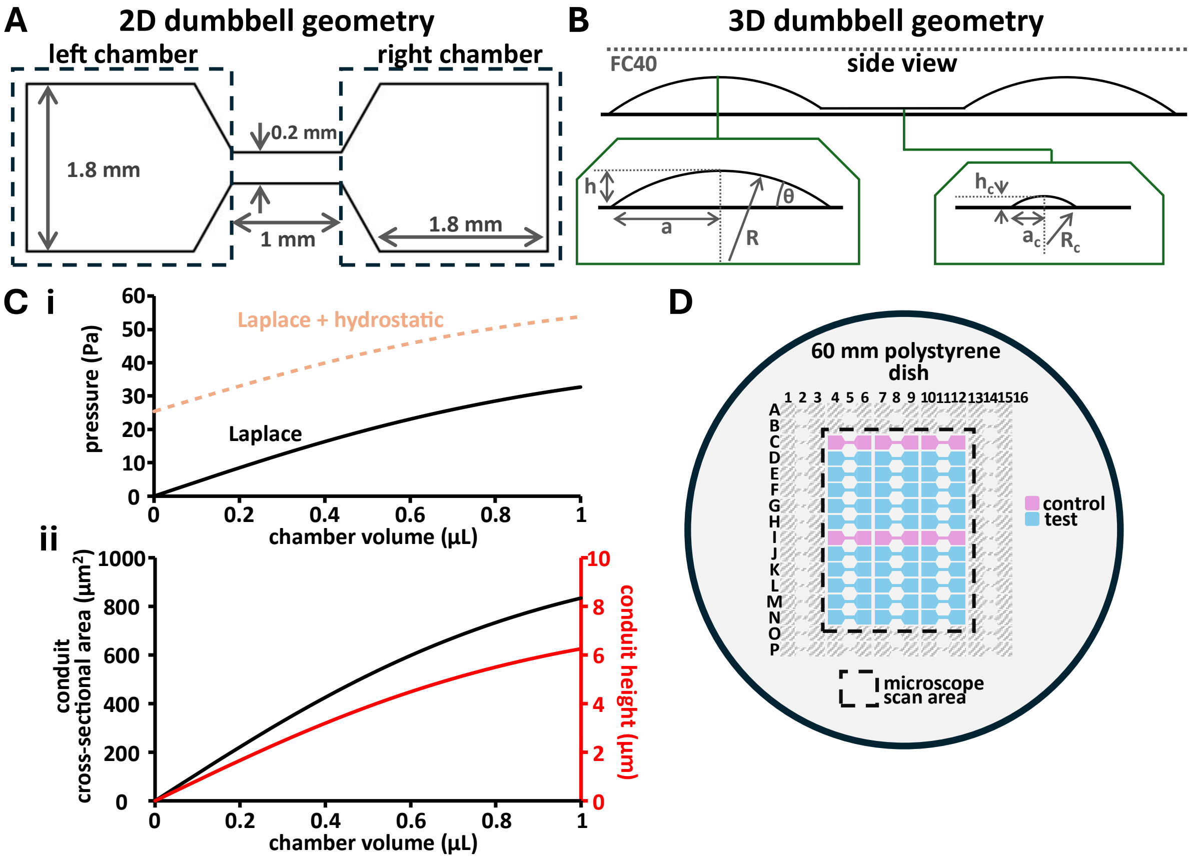


**Figure S2. Dumbbell geometry, pressures, and arrays. (A)** Single dumbbell with two chambers (1.8 mm x 1.8 mm) connected by a conduit (0.2 mm x 1 mm). **(B)** Side view of a dumbbell with equal volume in chambers. **(C)** Effect of increasing chamber volume on **(i)** chamber pressure, and **(ii)** conduit height and cross-sectional area. **(D)** Array of dumbbells on a 60 mm polystyrene dish. The central 36 dumbbells are within the area scanned by the Incucyte system (test dumbbells + 20% FBS are shown blue, and no-FBS controls pink).

### Evaluation of PBS washes for removal of residual FBS

In addition to minimising FBS concentration used in the initial fabrication of fluid walls, microenvironments were rinsed with PBS to further minimise any effect of any residual FBS on chemo-attractant concentration and hence migration. Microenvironments were made using RPMI + green-fluorescent BSA, at the equivalent concentration to 0.5% FBS (0.25 mg/ml BSA) and changes in fluorescent intensity used to assess wash efficiency (**Figure S3**). After collecting fluorescent images at low power, conduits connecting chambers are scarcely visible at this magnification, but green fluorescence is clearly visible in both dumbbells and the hexagonal spaces between conduits. A single PBS wash applied to dumbbells (but not hexagonal spaces) removed residual FBS from dumbbells (but not spaces); subsequent washes had minimal effect on dumbbell fluorescence.

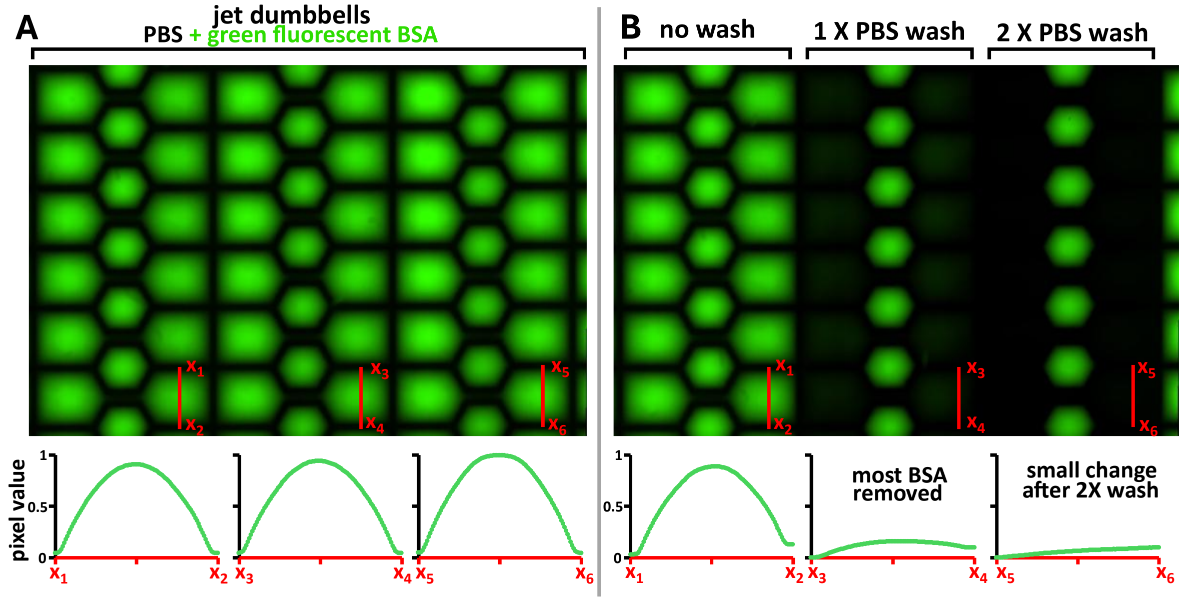


**Figure S3. Evaluation of PBS washes for removal of residual FBS. (A)** Microenvironments were made by first wetting a 60 mm dish with PBS + green-fluorescent BSA, withdrawing to leave a thin film, overlaying with FC40, and jet-printing dumbbells. Pixel intensities were measured across each chamber, averaged for left and right chambers across six dumbbells per column, and normalized to the maximum value. Intensity profiles are aligned to their corresponding column. **(B)** The middle column was washed once with 600 nl PBS per chamber, and the right washed twice. A single PBS wash removed most of the fluorescent signal, with the second providing only minor additional reduction.

#### Visualisation of flow between chambers

Flow between chambers was quantified through side-on imaging. Chamber heights were measured and volume estimated using spherical cap geometry, approximating each chamber as a round droplet with an equivalent footprint area. The flow rate was defined as the rate of change in volume, $Q = -dV/dt$, of the higher-volume chamber. To reduce noise, flow rate was calculated from chamber height after fitting a two-term exponential function in MATLAB to the temporal height profile of the larger chamber. Continuous flow was maintained during cell plating thereby confining cells to their respective chambers throughout the attachment phase.

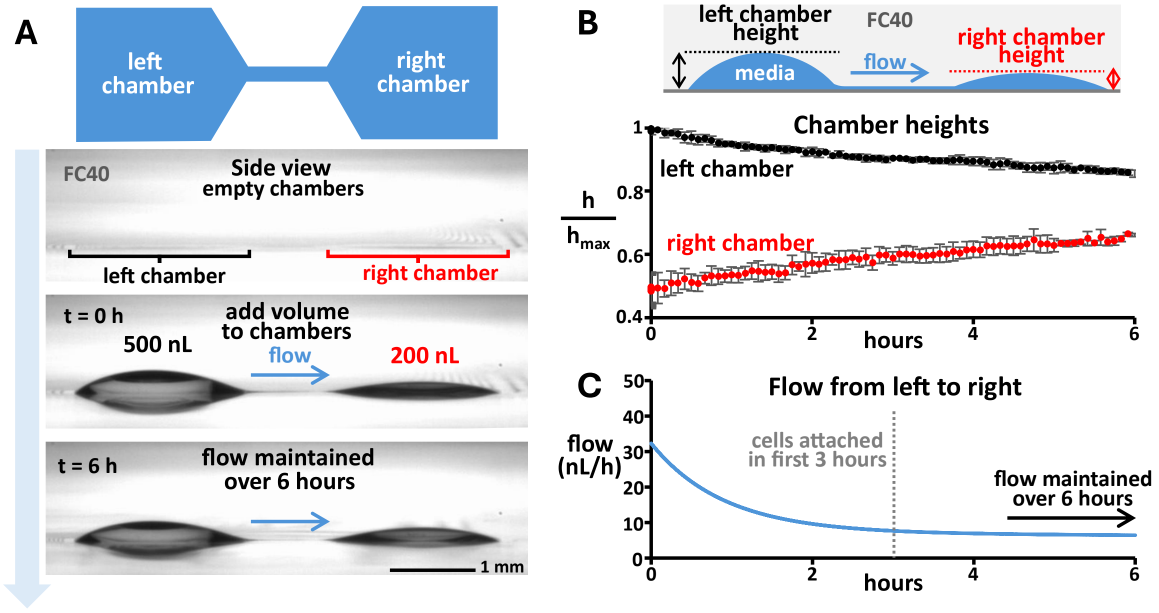


**Figure S4. Visualization of flow between chambers. (A)** Side-view imaging of a dumbbell follow withdrawal of residual media to leave empty chamber. Media volumes of 200 nl (right)and 500 nl (left) were added, establishing a pressure gradient driving flow from left to right. **(B)** Chamber heights were measured over time (± standard deviation, n = 3). **(C)** Approximating chambers as spherical caps enables flow-rate estimation. Flow during seeding confined suspended cells within chambers, with attachment occurring after ~3 h.

### Optimisation of cell seeding within dumbbells

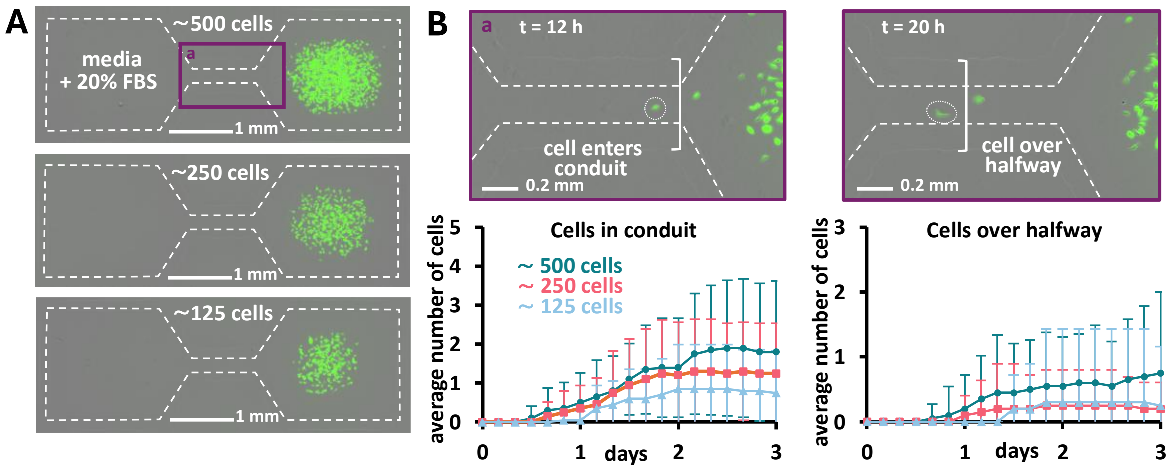


**Figure S5. Optimization of cell seeding within dumbbells. (A)** Cells were seeded into the right chamber at approximately 500, 250 and 125 cells per dumbbell. Media + 20% FBS was added to the left chamber. **(B)** Zoomed image of purple boxed region in A at two timepoints. The number of cells to enter the conduit and the number of cells over halfway into it for each condition (± standard deviation, 20 dumbbells in each condition).

### Diffusive transport of FBS within microenvironments

Determination of diffusion rates through microenvironments is essential in interpreting cellular response to chemical gradients. At equilibrium (no flow; equal chamber volumes), FBS diffuses from the left to the right chamber. The concentration profile along the conduit $c(x,t)$ was approximated using the one-dimensional diffusion equation derived from Fick's 2nd law for a semi-infinite medium and constant boundary concentration [1].

$$c(x,t) = C_o erfc\left(\frac{x}{2\sqrt{Dt}}\right) \quad \text{(S7)}$$

where $C_0$ is the starting concentration in the left chamber and $D$ is the diffusivity. Given that albumin is the principal component of FBS [2] we use green-fluorescent BSA as both a surrogate and a visual aid to confirm initial confinement to the left chamber. The diffusivity is approximated as that of albumin in water (61 μm$^2$/s [3]).

The time taken to reach steady state, $t_s$, was estimated as the time required for the concentration at the distal end of the conduit to reach 40% of the left chamber concentration – representing <5% difference between the concentration profile and linear approximation:

$$t_s = \frac{1}{D}\left(\frac{l}{2erfc^{-1}(0.4)}\right)^2 \quad \text{(S8)}$$

At steady state, the gradient is considered linear and stable for a time $t_{linear}$, defined as the time required for the right chamber to reach 5% of the left chamber concentration. Considering molecular transfer from the left to the right chamber:

$$t_{linear} = 1.05\frac{l m_{transfer}}{D A_c C_0} \quad \text{(S9)}$$

where $l$ is conduit length and $m_{transfer}$ the mass of molecules transferred from the left chamber to the right ($m_{transfer}$ = 1 μg, $A_c$ = 170 μm$^2$).

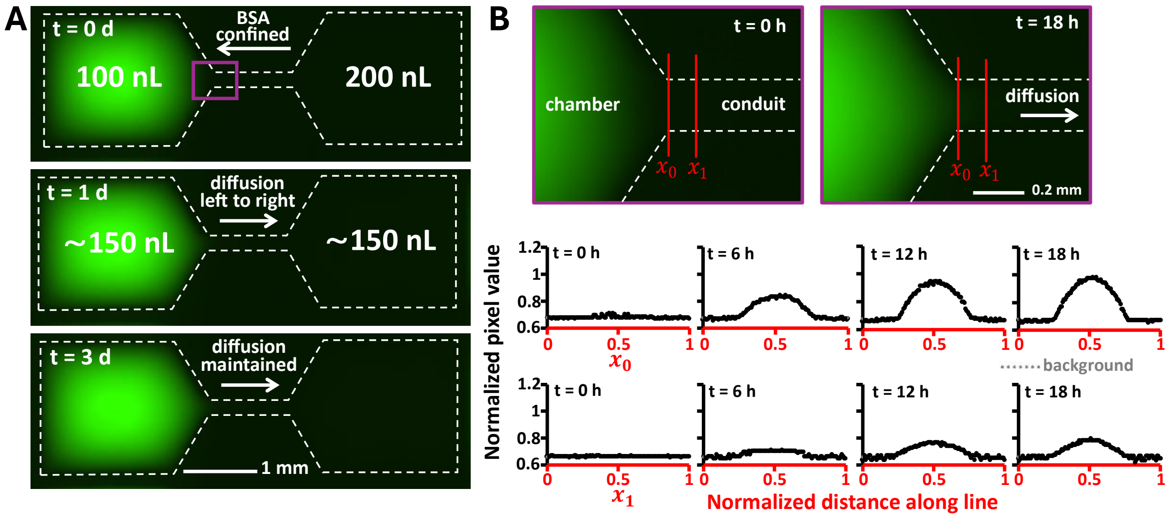


C

| | |
|---|---|
| Conduit length | 1 mm |
| Conduit cross sectional area at equilibrium | 170 μm$^2$ |
| Serum concentration in left chamber at equilibrium | 13.33% |
| Diffusivity | 61 μm$^2$/s |
| Time BSA first detected at conduit start | ~ 2 h |
| Time to establish stable diffusion gradient | ~ 3.5 h |
| Duration of stable diffusion gradient | ~ 9 d |

**Figure S6. Visualizing diffusion through dumbbells. (A)** Green-fluorescent BSA is used as a surrogate for FBS. Backflow driven by the gradient due to differences in Laplace pressures confines BSA initially to the left chamber. Chamber volumes equilibrate and BSA diffuses from left to right through the conduit. **(B)** Zoomed image of purple boxed region shown in A. Pixel intensity measured across the conduit opening $x_0$ (first row, 6 h intervals) and at 100 μm (second row, same timepoints) into the conduit, $x_1$, confirms diffusion of fluorescent BSA. Data is averaged across three reps, normalized to maximum pixel value and background-corrected using average pixel value along first 0.15 mm of line, outside of conduit **(C)** Summary of measured and calculated parameters. Diffusion is sustained for more than 3 d.

### Impact of chamber volume and serum concentration on cell migration and proliferation

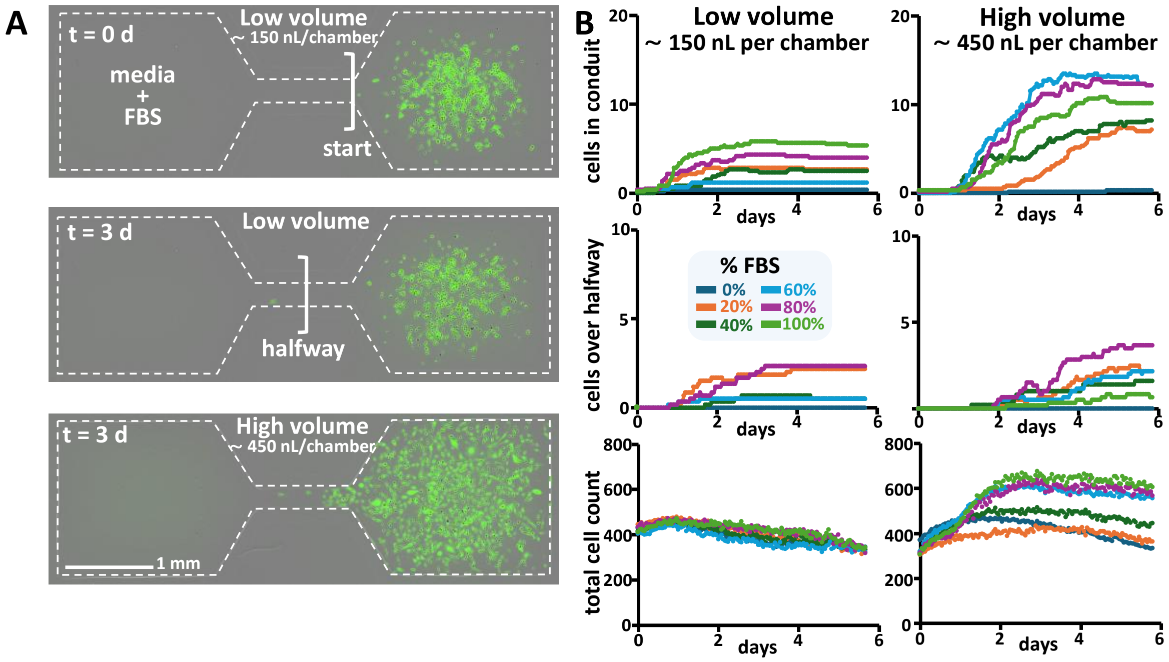


**Figure S7. Impact of chamber volume and serum concentration on cell migration and proliferation. (A)** Approximately 500 cells were seeded into the right chamber, with media containing varying concentrations of FBS added to the left chamber. In the *low-volume* condition, chamber volume at equilibrium was ~150 nl, and in *high-volume* there was ~450 nl per chamber. **(B)** Cells migrating into the conduit were counted at the entrance and at the halfway point for each serum concentration. Total cell numbers (cells within the conduit and chambers) were quantified to assess the effect of serum content and chamber volume on proliferation. Each line represents the average of 6 dumbbells.

### Optimizing cell picking

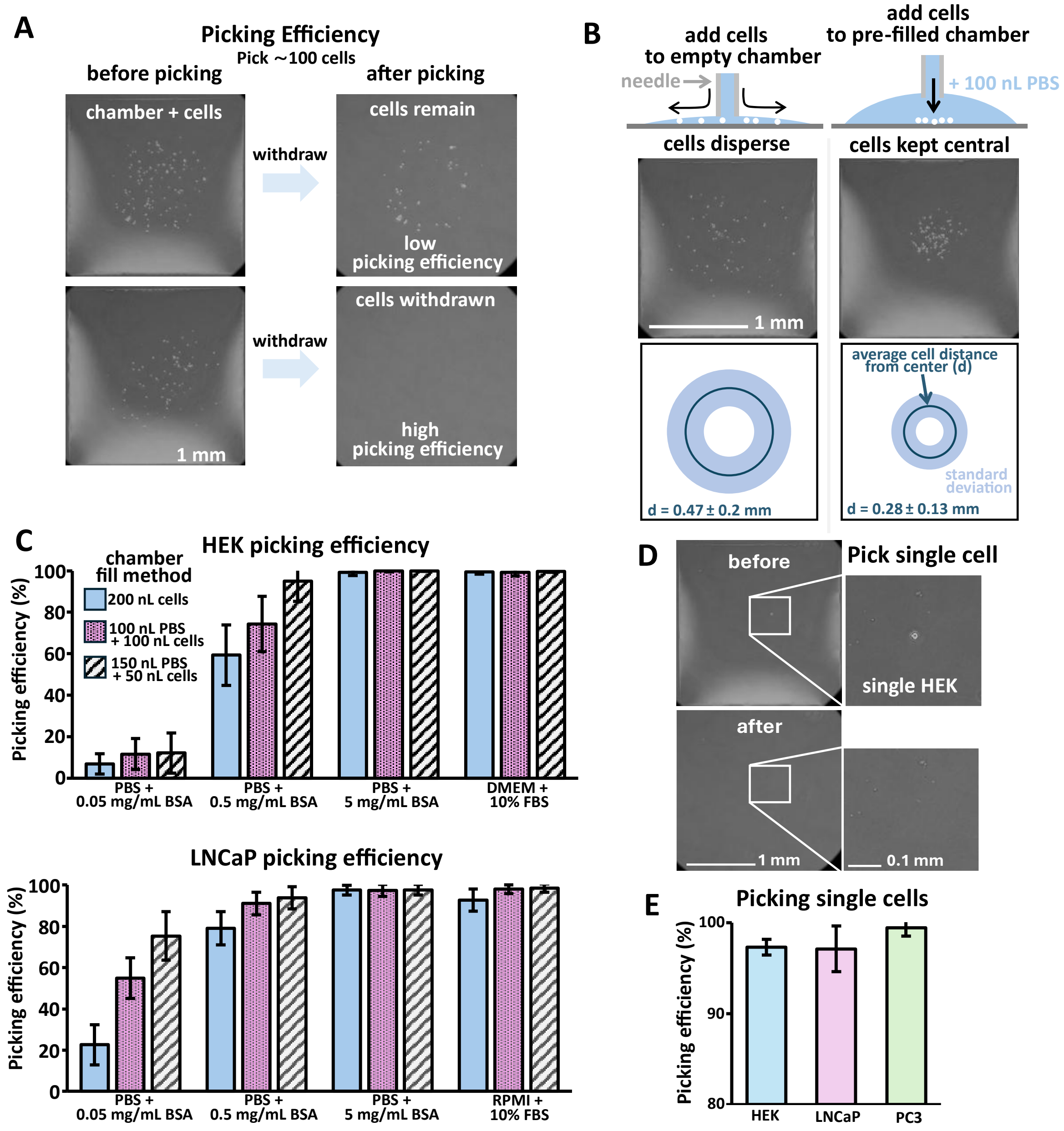


**Figure S8. Cell picking optimization. (A)** Examples of low and high picking efficiency. Approximately 100 cells were added to a chamber and the entire volume withdrawn. Picking efficiency was defined as the percentage of cells removed. **(B)** Seeding cells into empty chambers resulted in dispersal, whereas pre-filling with fluid kept cells centralized within the chamber (n = 10 chambers, each containing ~ 100 cells). **(C)** Picking efficiencies for HEK and LNCaP cells under varying conditions (± standard deviation, n = 3 technical replicates, picking from 50 grid chambers in each experiment). Chambers were fabricated with increasing BSA concentrations to inhibit adhesion, and the ratio of pre-fill chamber volume to added cell suspension was adjusted to maintain cell centrality and improve efficiency. **(D)** Example showing successful picking of a single cell. **(E)** Picking efficiencies for single HEK, LNCaP, and PC3 cells, using the optimized method (± standard deviation, n = 3).

### Automated cell segmentation and tracking using Cellpose and ImageJ

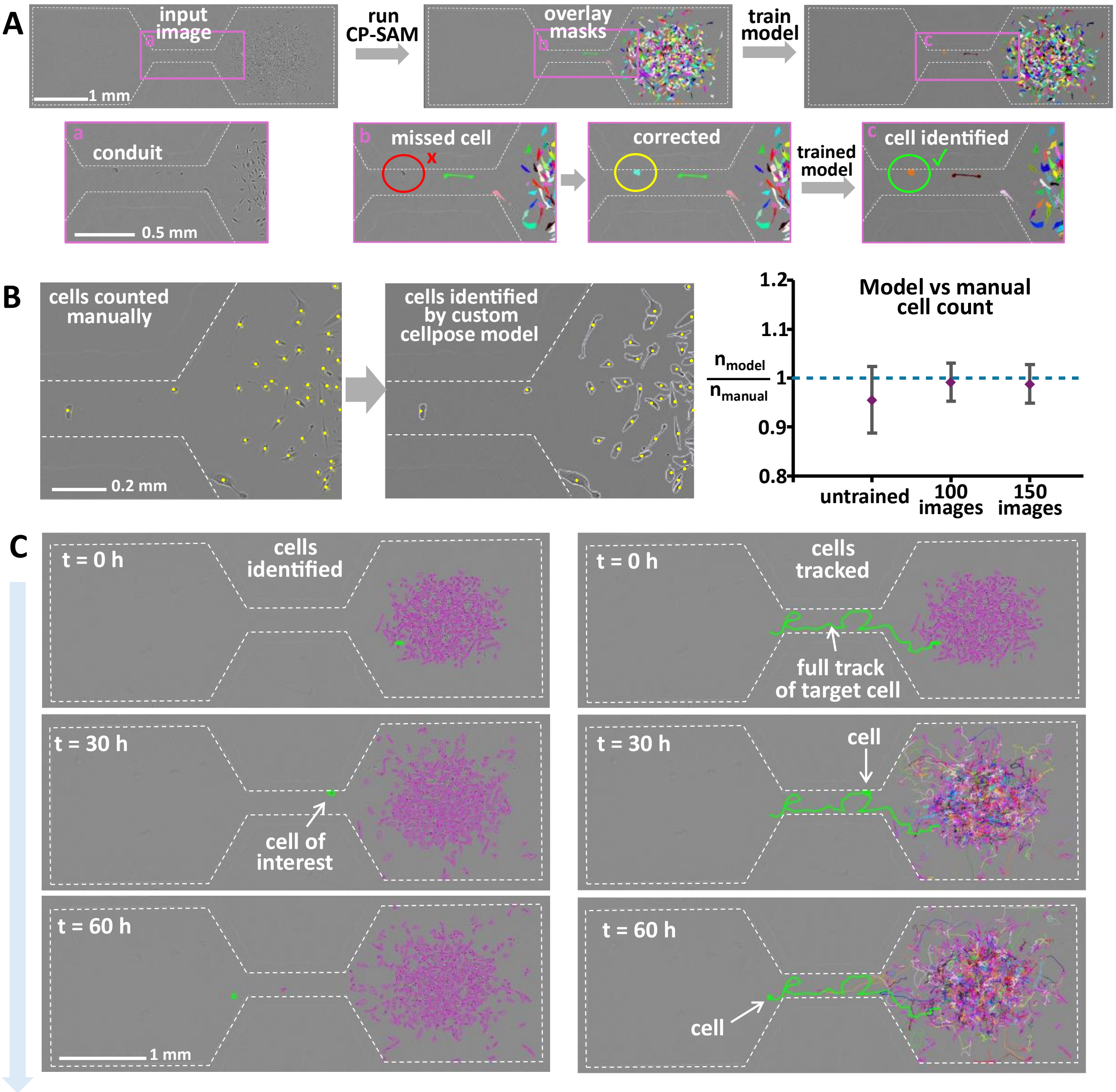


**Figure S9. Automated cell segmentation and tracking using Cellpose and ImageJ. (A)** Phase-contrast images of dumbbells were processed in Cellpose. Purple rectangles: zoom shown below. The CP-SAM algorithm was used for initial segmentation with missing cells corrected manually. The corrected mask is saved and the procedure repeated to train a custom model, which improved cell detection. **(B)** Agreement with manual cell counts was evaluated by comparing manual cell counts in 5 representative images with automated counts from the untrained model, after training on 100 images, and after training on 150 images (± standard deviation, n = 5). **(C)** The custom model was integrated with ImageJ to provide initial cell segmentation, followed by tracking to extract quantitative parameters of interest. Image contrast and brightness have been adjusted for clarity.

#### Descriptions of Supplementary Movies

**Movie S1**. Time-lapse movies of six representative dumbbells stitched together, each containing a migratory cell. In all cases, a single cell traverses the conduit into the opposing chamber, exhibiting diverse and dynamic morphological changes. Images were acquired hourly; the sequence is shown at increased speed and looped three times.

**Movie S2**. Magnified view of a single dumbbell showing pronounced morphological changes in the migrating cell over time. Images were acquired and presented as described above.

**Movie S3**. Magnified view of single dumbbell. In this example, the migrating cell appears more motile, with small morphological changes compared to Movie S2. Images were acquired and presented as described above.

**Movie S4**. Three example dumbbells where Cellpose was used for cell segmentation. In each case, the migratory cell of interest is highlighted in green. Dumbbells are stitched together from three experiments.

**Movie S5**. Cell tracks overlaid following segmentation and tracking in ImageJ. The same three dumbbells shown in Movie S4 are displayed, with the trajectory of the migratory cell highlighted in green.

**Movie S6**. Heterogeneity within the migratory subpopulation revealed by cluster analysis, which identified two groups: smaller, faster-moving cells and larger, slower-moving ones. A representative example from each group is shown side-by-side for comparison.

#### Supplementary Information References

[1] F. Nebuloni, Q. B. Do, P. R. Cook, E. J. Walsh, and R. Wade-Martins, ‘A fluid-walled microfluidic platform for human neuron microcircuits and directed axotomy’, *Lab on a Chip*, vol. 24, no. 13, pp. 3252–3264, Jun. 2024, doi: 10.1039/D4LC00107A.
[2] G. L. Francis, ‘Albumin and mammalian cell culture: implications for biotechnology applications’, *Cytotechnology*, vol. 62, no. 1, pp. 1–16, Jan. 2010, doi: 10.1007/s10616-010-9263-3.
[3] A. K. Gaigalas, J. B. Hubbard, M. McCurley, and S. Woo, ‘Diffusion of bovine serum albumin in aqueous solutions’, *J. Phys. Chem.*, vol. 96, no. 5, pp. 2355–2359, Mar. 1992, doi: 10.1021/j100184a063.
